# Unbiased genetics identifies a glutamatergic signaling network as a mediator of daily sleep patterns

**DOI:** 10.1101/2025.05.08.652901

**Authors:** Sawyer Grizzard, Julia S. Lord, Alicia L. Ravens, Yajing Hao, Wei Tang, Sung Hyun Lee, Sarah Jo Sleiman, Kathryn K. Joyce, Kevin D. Donohue, Bruce F. O’Hara, Fei Zou, Tal Kafri, Graham H. Diering

## Abstract

Sleep is a fundamental, conserved behavior important for survival. In many species, sleep behavior is controlled by poorly understood interactions between a system of circadian rhythms (CR), which promote sleep at ecologically appropriate times, and a homeostatic sleep drive that accumulates with time awake. The CR is a cellular phenomenon, driven by molecular oscillations of “clock genes” in nearly all cells. Emerging evidence indicates neuronal synapses are a key locus for the accumulation and resolution of sleep need, supporting a cellular basis of sleep need. Indeed, efforts to understand the genetic basis of sleep need identified Homer1a, a regulator of synapse homeostasis. To develop further insight into the genetic basis of sleep regulation, we measured daily sleep patterns in genetically diverse strains of mice from the Collaborative Cross. Strains with 1) highly consolidated light-phase sleep, or 2) fragmented, arrhythmic sleep, were identified for genetic analysis using quantitative trait loci (QTL) mapping. Excitingly, in F2 hybrids, 19 of 32 metrics of sleep and circadian behavior mapped to a narrow QTL containing GRM5, a postsynaptic glutamate receptor and binding partner of Homer1a, and GCPII, an astrocytic enzyme that regulates NAAG, a peptide agonist for the presynaptic/astrocytic glutamate receptor GRM3. Collectively, these genes form a coordinated glutamatergic signaling network across the tri-partite synapse. Pharmacology targeting GRM5, GCPII, and GRM3 strongly modulated sleep, functionally validating them as sleep-regulating genes. Our findings support a model in which synapses act as a cellular site for integration of circadian and sleep-need signals to regulate daily sleep patterns.

**Significance Statement:** Sleep is a fundamental pillar of human health. Restorative sleep is supported by circadian rhythms (CR). Sleep and CR disruptions are prevalent in neuropsychiatric conditions and are increasingly understood as causal drivers of altered brain function and behavior. Therefore, sleep and CR are attractive therapeutic targets to improve health. However, the basic biological mechanisms that couple our need to sleep with the CR are not understood. Genetic discovery using diverse strains of mice identifies a glutamatergic signaling network coordinated across the synapse as an underlying genetic basis for daily sleep patterns. This work builds upon a growing body showing that excitatory synapses are a cellular site for the resolution of sleep need and interaction with the CR.

## Introduction

The need for sleep is conserved across animal species, accumulates during wake and can only be dissipated by sleeping^1^. The basis for the restorative functions of sleep, that resolve sleep-need, are poorly understood^2^. In most species, the timing and quality of sleep are understood to be controlled by the interaction of two major factors: a circadian rhythm (CR) component (called process C in the two-process model), that promotes sleep at the ecologically appropriate time of day, and the homeostatic sleep drive (process S), that promotes sleep in proportion to time spent awake^3^. The CR is understood as a cellular phenomenon, based on functionally interacting clock genes within most cells of the body^4^. In contrast, the molecular basis of the homeostatic sleep drive is largely unknown. Furthermore, although the homeostatic sleep drive and CR have largely been understood as independent mechanisms, mounting evidence suggests that these processes functionally interact^5-7^, although the cellular or molecular basis of this interaction remains unknown.

In mammals, slow-wave activity (SWA), also called delta-power, measured by electroencephalography (EEG) during non-rapid eye movement (NREM) sleep, increases as a function of time spent awake and dissipates with time asleep, establishing NREM-SWA as a physiological marker for the accumulation and resolution of sleep-need^2,5^. NREM-SWA accumulation varies between mouse strains and is genetically determined, indicating that EEG-defined sleep-need has a molecular substrate^8^. Subsequent efforts identified Homer1a, encoding a synaptic protein, as a molecular correlate of sleep-need^9,10^. Homer1a is an immediate early gene induced in neurons in response to heightened activity and subsequently drives homeostatic scaling-down, a global weakening of excitatory synapses^11^, through interaction with type-I metabotropic glutamate receptors mGluR1/5^12^ (encoded by GRM1 and GRM5). Furthermore, Homer1a is potently induced in the mammalian forebrain in response to sleep deprivation (SD)^9,10^, and drives homeostatic scaling-down of excitatory synapses during sleep through interaction with mGluRs^13^. Thus, identification of homeostatic scaling factor Homer1a as a molecular mediator of sleep need clearly implicates neuronal synapse homeostasis in the restorative basis of sleep function. These findings strongly support the synapse homeostasis hypothesis (SHY), a long-standing model describing the synaptic basis for the benefits of sleep. SHY posits that neuronal synapses are strengthened during wake to encode information from experience, while during sleep, synapses are broadly weakened to restore synapse homeostasis^2^. Recent advances further show that the accumulation and resolution of sleep need involve intracellular kinase/phosphatase signaling acting in part on synapse proteins, supporting a view that sleep-need has a biochemical basis centered on synapse biology^14-17^.

Understanding of the molecular mechanisms for the timing and need for sleep has been enabled through unbiased genetic discovery in model organisms including mice^18^. Forward-genetics screens based on random mutagenesis in mice successfully identified functionally important genes, such as the sleep-promoting kinase SIK3 and its substrate HDAC4/5^19-21^. However, many sleep traits are believed to be highly polygenic and therefore dependent on genetic background^18^. Genetic reference populations provide a powerful system to study genetically complex traits. The collaborative cross (CC) is a collection of recombinant inbred mouse strains derived from systematic crossbreeding of eight progenitor strains: A/J, C57BL/6J, 129S1/SvImJ, NOD/LtJ, NZO/HlLtJ, CAST/EiJ, PWK/PhJ, and WSB/EiJ. The CC contains a high degree of genetic diversity comprising more than 36 million single-nucleotide polymorphisms^22,23^. Thus, the CC is ideal to advance understanding of genetically complex traits through unique allele combinations inherited from genetically well-characterized founder strains.

Here we screened sleep and CR phenotypes in a subset of CC strains, identifying strains CC036 with highly consolidated light phase sleep, and CC057 with highly fragmented sleep lacking a robust daily rhythm. Subsequent generation of F2 hybrids enabled quantitative trait loci (QTL) mapping. Using a non-invasive piezoelectric based home-cage monitoring system, PiezoSleep, we examined 32 metrics of sleep and circadian behavior. Excitingly,19 out of 32 metrics, primarily related to the consolidation and timing of sleep, mapped to a strong and narrow QTL on chromosome 7 containing glutamate carboxy peptidase II (GCPII, also called FOLH1) and GRM5. GCPII is a cell-surface enzyme primarily expressed on astrocytes that degrades an abundant neuroactive peptide N-acetyl aspartyl glutamate (NAAG), a known agonist for metabotropic glutamate receptor GRM3, expressed on neuronal pre-synapses and astrocytes^24-28^. Collectively, GRM5, GCPII, and GRM3 form a glutamatergic signaling network coordinated across pre- and post-synaptic compartments and astrocytic processes, called the tri-partite synapse. Pharmacology targeting these proteins significantly affected sleep behavior in the parental CC strains, supporting our identification of functionally important sleep genes. These findings, like the identification of Homer1A in prior studies, reinforce the role of synapse biology in sleep regulation. Given that GRM5 is a known binding partner of Homer1A^12,13,29^, our unbiased discovery approach suggests that synapses may serve as a convergence point of circadian and homeostatic sleep regulation.

## Results

### Daily sleep phenotypes in genetically diverse strains of mice from the collaborative cross

To begin our investigation of how genetic variation shapes sleep phenotypes across the 24hr day, we analyzed daily sleep behavior patterns across a panel of 13 strains of mice from the Collaborative Cross (CC), including both sexes, in comparison to well-characterized C57Bl/6J (B6). The CC collection is derived from the systematic interbreeding of eight genetically diverse founder lines, followed subsequently by inbreeding to generate recombinant inbred strains^22^ (Fig1A). Following a period of acclimation, daily sleep patterns were recorded uninterrupted over 7 days on a 12:12hr light/dark cycle, using a non-invasive, home-cage sleep recording system, PiezoSleep. This system uses highly sensitive piezoelectric sensors to quantify mouse motions and breathing to score wake/sleep behavior and has been validated using simultaneous EEG and live-video scoring^30-32^. Continuous hourly sleep traces over multiple days revealed striking variations in the daily sleep patterns between mouse strains (Fig.1B). We noted particular differences between two strains, CC036 and CC057, respectively showing highly consolidated sleep in the light phase and robust daily rhythms, or fragmented sleep distributed over the light/dark cycle and lacking a robust daily rhythm (Fig.1B). We then quantified average daily sleep amount and bout lengths across 24hrs or 12hr light/dark phases (Fig.1C-G); CC strains were compared statistically to B6, which should not be considered a control, but rather a benchmark from a well characterized genetic background. Average 24hr data revealed a range of total sleep amount and bout lengths, a metric of sleep consolidation. Interestingly, B6 mice exhibited lower amounts of total daily sleep, yet they showed the highest average sleep bout lengths; both measures significantly differed from several CC strains, including CC057, which had particularly short and fragmented sleep bouts (Fig. 1C-D). Average 24hr hourly traces show variable distributions of sleep across the light/dark cycle, showing that CC036 and CC057 sleep more than B6 in the light or dark phases respectively (Fig.1E-F). As expected for a nocturnal species, all 14 strains have higher sleep amount and bout lengths in the light phase, with CC036 showing the highest levels of total light phase sleep; however, CC057 showed a notably weaker separation of sleep amount/bout length between light and dark (Fig.1F-G). We then calculated the light/dark ratio for total sleep amount and bout length, which combined provides an overview of the degree of light phase consolidation, found to be highly divergent across the 14 strains (Fig.1H). Interestingly, CC036 and CC057 were found at opposite ends of this daily consolidation spectrum, with B6 being intermediate. Overall, divergent phenotypes suggest that innate genetic variation drives distinct daily sleep behaviors. To advance this idea, we selected CC036 and CC057 for subsequent genetic analysis using QTL mapping.

**Fig. 1.**
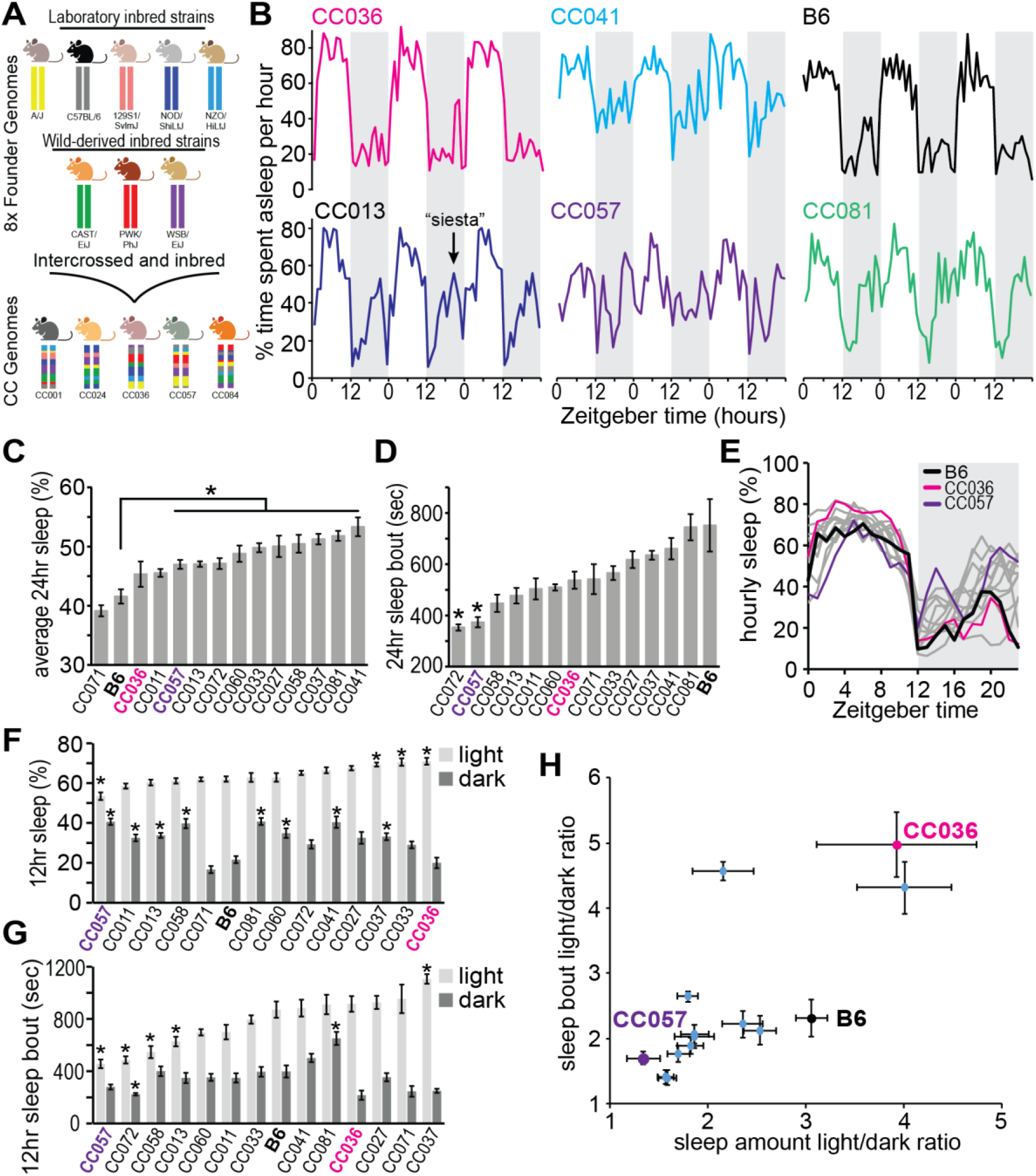
Daily sleep phenotypes in genetically diverse strains of mice from the collaborative cross. **A**. The collaborative cross (CC) is a collection of genetically diverse inbred mouse lines generated from systematic cross breeding of 8 founder mouse strains. **B-H**. Sleep behavior recorded continuously for 7days from 13 CC strains and C57Bl6/J (B6). Data averaged from 7-10 individuals per strain, including both sexes. Error bars indicate standard error. Statistical difference from B6 indicated, *P<0.05 Unpaired Student’s t-test with Bonferroni correction. **B**. Example 72hr hourly sleep traces. Dark phase indicated by shaded area. **C**. 24hr average time asleep (%). **D**. 24hr average bout length (sec). **E**. Overlaid 24hr hourly sleep % traces, averaged from 7 days, indicate variability in daily sleep patterns among 14 mouse strains. B6, CC036, and CC057 are indicated. Dark phase indicated by shaded area. **F**. Averaged sleep % for light and dark phases. CC057 and CC036 show the lowest and highest light phase sleep amount respectively. **G**. Average sleep bout length (sec) for light and dark phase. **H**. Scatterplot of light/dark ratios for sleep amount and sleep bout lengths. Upper right quadrant indicates strongest consolidation of daily sleep in the light phase. Note that CC036 shows the strongest consolidation of light phase sleep, CC057 shows the weakest consolidation, and B6 shows an intermediate level.

### Convergence of CC036 and CC057 sleep traits in the F1 hybrid

Next, we crossed CC036 and CC057 to generate F1 and subsequent F2 hybrids. Because this effort would involve hundreds of mice, we conducted all following analysis using only female mice to mitigate anticipated issues with aggression. To explore how allelic interactions shape sleep, we assessed sleep in F1 hybrids in comparison to parental CC036 and CC057 using our non-invasive home-cage system (Fig.2A). In multi-day sleep traces, F1 hybrids showed qualitative features of both parents, and in most quantitative metrics showed convergent daily sleep phenotypes. F1 hybrids showed a prominent delay in sleep onset after lights-on, similar to CC057 (Fig. 2A-B), whereas, unlike CC057, they showed a more prominent consolidation of sleep amount and bout lengths in the light phase and higher light/dark sleep amount ratios (Fig.2B-D). We then compared the distribution of sleep or wake into different bout lengths, a measure of consolidation of wake/sleep behavior. Consistent with our earlier recording (Fig.1D), CC057 shows highly fragmented sleep with a clear shift towards short bout lengths, whereas CC036 showed comparably consolidated sleep of longer bout lengths. In this metric, F1 hybrids were intermediate between parental strains (Fig.2E), consistent with their polygenic inheritance. CC057 also showed fragmented wake: more wake episodes of short duration and low levels of consolidated wake of longer durations. Interestingly, F1 hybrids expressed longer episodes of consolidated wake compared to parental strains (Fig.2F), perhaps an emergent phenotype of hybridization. Next, we used our non-invasive system to estimate rapid eye movement (REM) and NREM sleep composition in parental and F1 hybrid strains. Sleep scoring using PiezoSleep is based on the detection of breath rate, the physical detection of diaphragm and chest movement of the sleeping mouse^30,32^. NREM and REM can be estimated based on rapid changes in breath rate and regularity accompanying state transitions^30,33^. Estimated NREM was found to be significantly different in daily patterns and consolidation between CC057 and CC036, and again, F1 hybrids showed a convergence of these traits (Fig.S1A-D). In contrast, estimated REM showed no or minimal differences (Fig.S1E-H). Overall, the convergence of daily sleep phenotypes in F1 hybrids is consistent with the polygenic nature of daily sleep behavior and supports the existence of divergent alleles that significantly influence sleep traits in the parental strains.

**Fig. 2.**
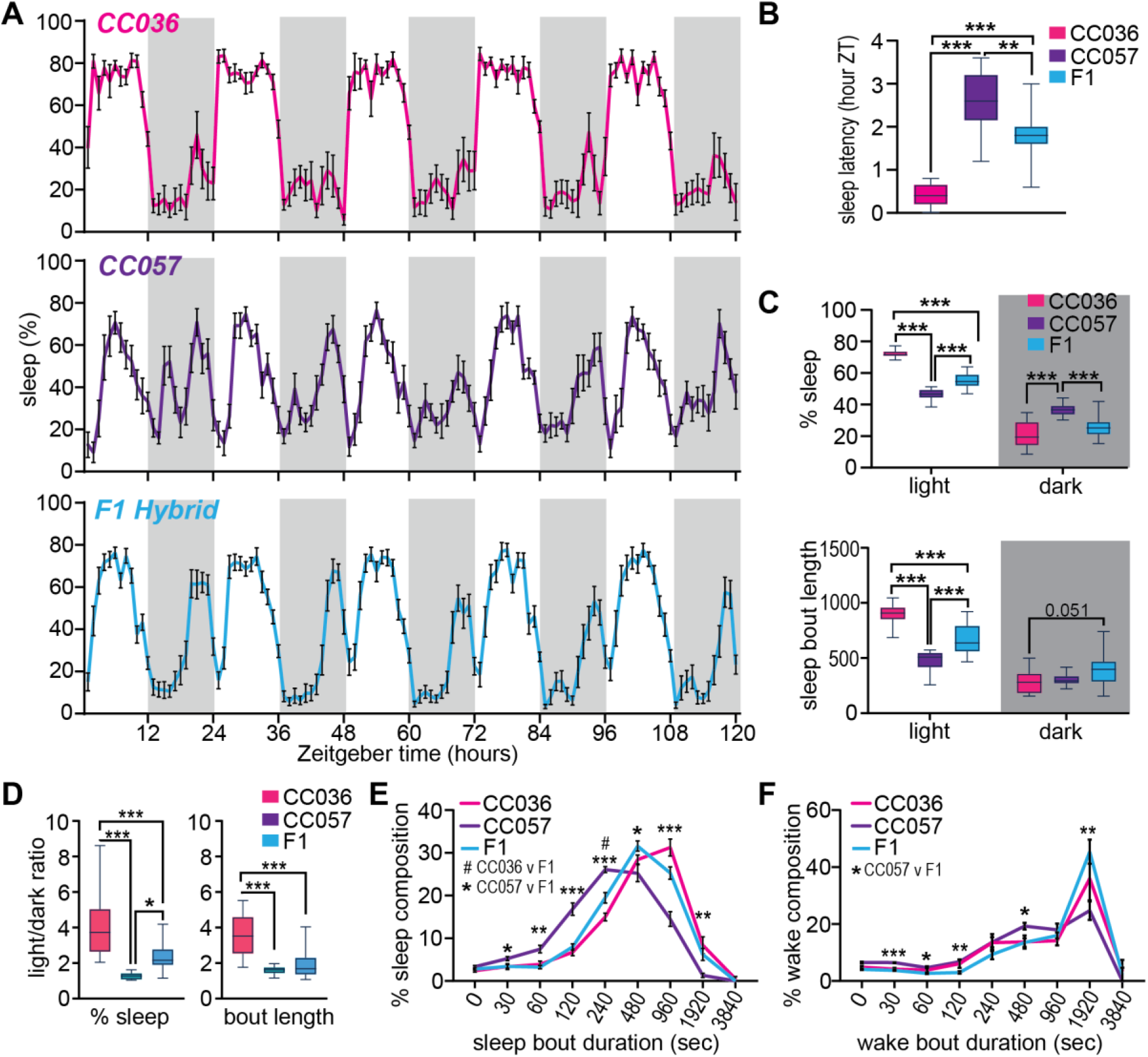
Convergence of CC036 and CC057 sleep traits in the F1 hybrid. **A**. Trace of average sleep % per hour over 5 days from female CC036 and CC057 parental strains and in the F1 hybrids. Dark phase indicated by shaded area. Error bars indicate standard error, N=10 females each for CC036 and CC057, and N=20 for F1 females. **B**. Average latency to sleep after light onset. **C**. Quantification of average % sleep or bout length (sec) in light and dark phases. **D**. Light/dark ratios of % sleep and bout length for CC036, CC057, and F1 hybrids. *P<0.05, **P<0.01, ***P<0.001 1-way ANOVA with Tukey’s multiple comparison test. F1 hybrids show intermediate phenotypes in most metrics between parental CC036 and CC057. **E-F**. Cumulative histogram of daily distribution of sleep (E) or wake (F) into bout lengths of varied duration. CC057 show distribution of wake and sleep into shorter bouts indicating overall fragmentation of wake/sleep activity. F1 hybrids show intermediate distribution of sleep bout lengths, and consolidation of wake into longer bouts. *P<0.05, **P<0.01, ***P<0.001 2-way ANOVA with Tukey’s multiple comparison test.

### Divergence of daily sleep traits in F2 hybrids

Next, we generated F2 hybrids; daily sleep patterns from 279 F2 females were analyzed over 5 uninterrupted days using our non-invasive home cage system. We then quantified 32 metrics of sleep and circadian behavior from each F2 individual for subsequent QTL mapping (summarized in Fig.S2). This included metrics of total sleep and wake, estimated REM and NREM, and circadian metrics: cosinor amplitude and acrophase, fast Fourier transform (FFT), and daily correlation coefficients, measures of the robustness of daily rhythms^34-36^. Multi-day hourly sleep traces revealed a continuum of phenotypes, from highly consolidated CC036-like sleep to fragmented, arhythmic CC057-like patterns, with many intermediate variants (Fig.3A). Circadian analysis showed similar variability: while some mice exhibited strong, regular rhythms, others displayed disrupted sleep-wake cycles. As examples, we highlight F2 individuals #90 and #97 as representative F2s with CC036 or CC057-like traits, respectively (Fig.3B-D). Plotting the light/dark ratios of sleep percentage and bout length across the F2 population, further emphasized this phenotypic diversity that exceeded the phenotypic range seen in CC036 and CC057 parental strains (Fig. 3E). Rank-ordered trait distributions showed a continuous spread for sleep traits rather than discrete clusters (Fig.3E-F, and Fig.S2), consistent with a polygenic basis of daily sleep phenotypes. Collectively, divergence of daily sleep phenotypes in the F2 generation points to a strong genetic and polygenic basis for variation in daily sleep patterns and supports the presence of divergent genetic alleles, inherited from the parental strains, that powerfully affect sleep phenotypes in this F2 population.

**Fig. 3.**
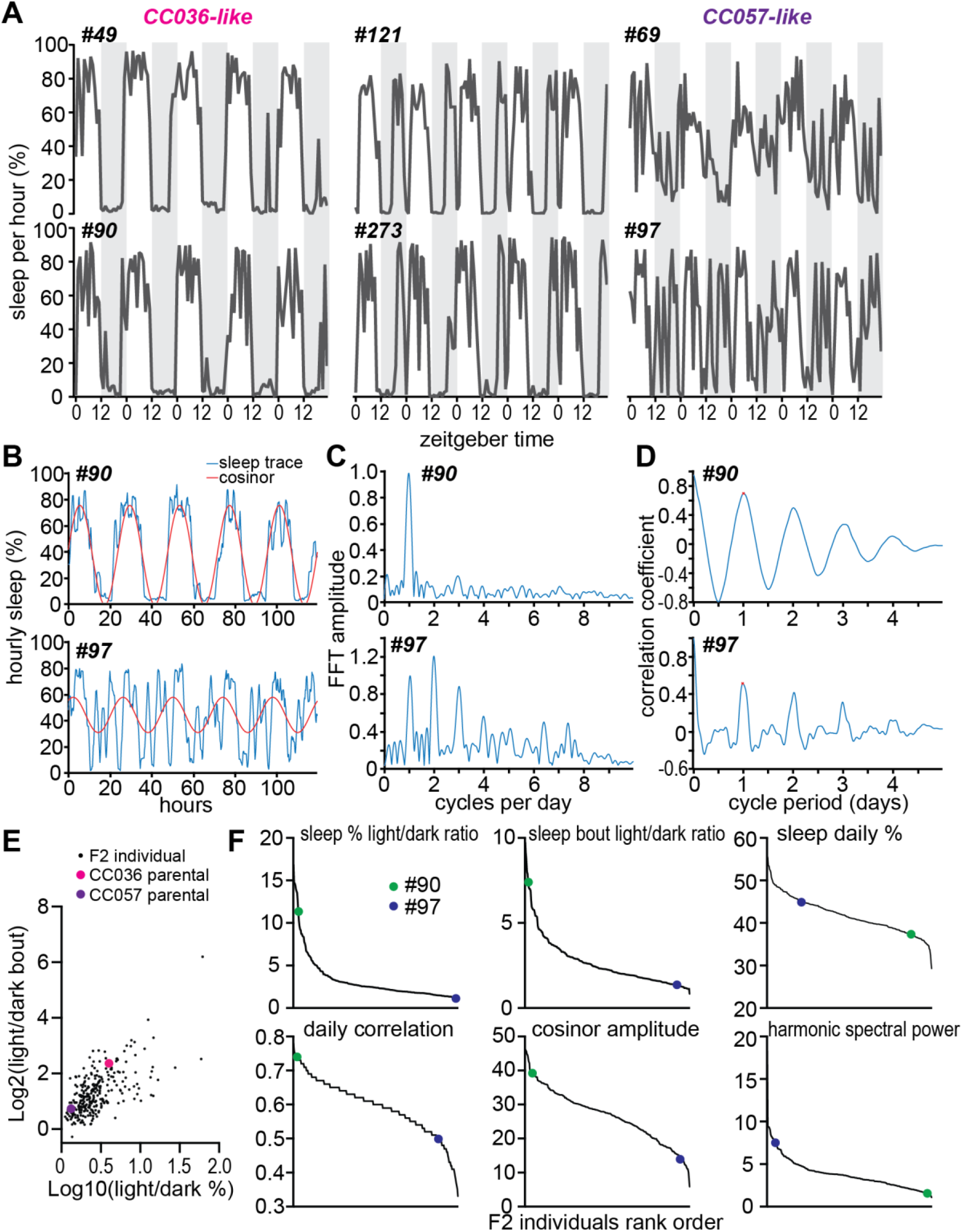
Divergence of daily sleep traits in F2 hybrids. **A**. Example 5-day hourly sleep traces for individual female F2 hybrids, dark phase indicated by shaded area. Note that some individuals in the F2 generation resemble the parental CC036 or CC057 daily sleep patterns. **B-D**. Examples of circadian analysis for F2 individuals #90 and #97, representing CC036 and CC057-like traits respectively. **B**. Cosinor analysis was conducted for 5day sleep traces. **C**. 5day sleep traces underwent fast Fourier transform (FFT) to determine harmonic spectral power. **D**. Daily correlation analysis of 5day sleep traces. **E**. Scatterplot of light/dark ratios for sleep amount (%) and sleep bout lengths for 279 F2 individuals. Average values from female CC036 and CC057 parental strains are indicated for reference. **F**. Population data obtained from 279 F2 individuals, showing ranked order highest to lowest values for indicated sleep and circadian metrics. For reference, individuals #90 and #97 are indicated.

### QTL mapping implicates a glutamate signaling network as a regulator of daily sleep

Parental CC036 and CC057, and F2 individuals were genotyped for single-nucleotide polymorphisms. Using 32 metrics obtained from F2 individuals (Fig.S2) we next performed a genome-wide QTL analysis. Multiple QTLs reached genome-wide significance (Fig.S3 and TableS1), consistent with the polygenic nature of sleep behavior. Incredibly, 19 of 32 phenotypic traits resulted in an overlapping, highly prominent QTL located on chromosome 7, that exhibited elevated logarithm of odds (LOD) scores, including %sleep light/dark ratio and cosinor amplitude (Fig.4A, Fig.S3, and TableS1). QTLs with lower LOD scores were also detected for a subset of traits on chromosomes 1, 6, 9, 15, and 17 (Fig.S3 and TableS1). Weaker, nonoverlapping QTLs that did not cross the 95% confidence interval were detected for estimated REM bout length in the light or dark phases on chromosomes 15 and 5 respectively (Fig.S3). While each quantitative trait enabled genome-wide statistical analysis, several traits are not independent processes. Indeed, Spearman correlation revealed a high degree of correlation amongst total sleep, wake, estimated NREM, and circadian metrics, but not estimated REM metrics (Fig.4B). A shared prominent QTL among 19 different quantitative phenotypic traits suggests a shared underlying genetic mechanism. The minimal genomic overlap among these 19 traits was localized to a 4.83 megabase region of chromosome 7 between 84.49-89.31MB (TableS1). In numerous instances, neighboring genes function in shared biological processes^37^. Excitingly, this region contains two glutamatergic synapse-related genes GRM5 and GCPII (glutamate carboxypeptidase II, also called FOLH1)(Fig.4C). GRM5 encodes the postsynaptic metabotropic glutamate receptor 5 (mGluR5), which interacts with Homer1a, a known molecular marker of sleep need^9,10^. Moreover, GRM5 has been implicated in sleep biology across numerous studies^13,38-41^. GCPII encodes an astrocytic cell-surface enzyme that degrades an abundant neuroactive peptide N-acetyl-aspartyl-glutamate (NAAG), a known selective agonist for presynaptic metabotropic glutamate receptor 3^24-27^ (mGluR3, encoded by GRM3). GRM3 is localized to excitatory pre-synapses and is the primary mGluR expressed in adult astrocytes^28,42,43^. Thus, QTL mapping implicates a glutamatergic signaling network coordinated across the tri-partite synapse as a regulator of daily sleep patterns.

**Fig. 4.**
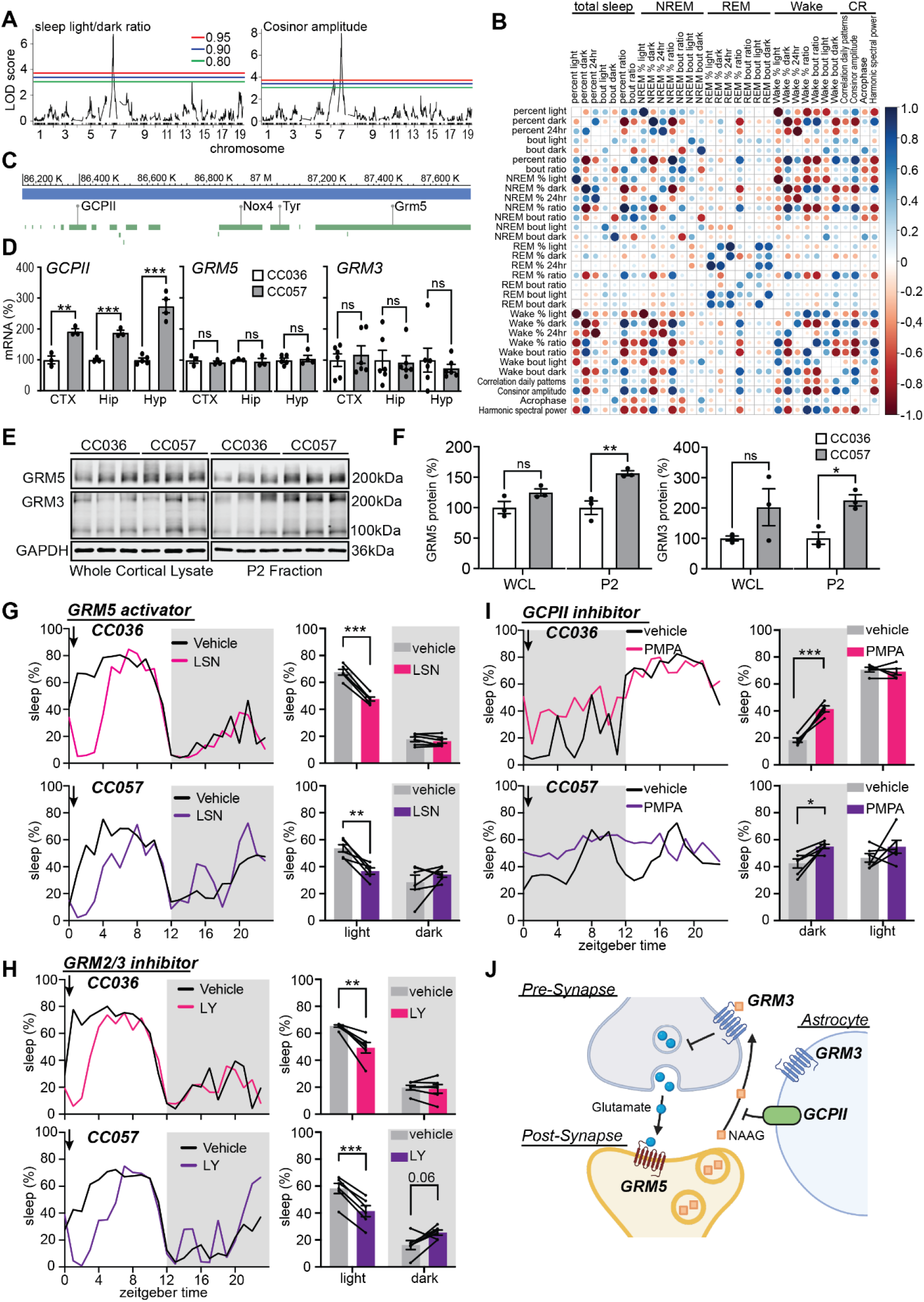
QTL mapping identifies a glutamate signaling network as a regulator of daily sleep. **A**. Significant genome-wide QTLs, indicated by logarithm of odds (LOD) score, associated with sleep patterns in F2 mice were detected for sleep and circadian metrics, examples shown for %sleep light/dark ratio and cosinor amplitude. 95%, 90%, and 80% significance levels indicated: red, blue, green lines respectively. **B**. Spearman correlation between sleep and circadian metrics. The sizes of circles show the absolute value of corresponding correlation coefficients. Color indicates the value of correlation coefficients, positive in blue, negative in red. **C**. Map of chromosome 7 region containing candidate genes GRM5 and GCPII. **D**. qPCR measurement of GCPII, GRM5, and GRM3 mRNA isolated from cortex (CTX), hippocampus (Hip), and hypothalamus (Hyp) from CC036 and CC057 mice, values normalized to CC036. **P<0.01, ***P<0.001, ns: not significant, Unpaired Student’s t-test. N=3-6. **E-F**. Western blot of GRM5 and GRM3 from whole cortical lysate (WCL) or synapse enriched P2 fractions from CC036 and CC057 mice. Quantification of western blots shows trend in WCL and significantly increased expression in P2 of GRM5 and GRM3 in CC057 compared to CC036. *P<0.05, **P<0.01, ns: not significant, Unpaired Student’s t-test. N=3. **G-I**. Sleep recordings and pharmacology validation of QTL hits. CC036 and CC057 mice were injected with vehicle followed 72hrs later by drug, total sleep amount (%) was compared between vehicle and drug treatment days. Time of injection indicated by arrow. Mice injected at light onset with GRM5 activator LSN2814617 (LSN) (**G**), or GRM2/3 inhibitor LY341495 (LY) (**H**), or dark onset with GCPII inhibitor 2-PMPA (PMPA) (**I**). *P<0.05, **P<0.01, ***P<0.001 Paired Student’s t-test. N=5-6 per condition. **J**. Model depicting coordinated signaling of GRM5, GCPII and GRM3 across the tripartite synapse composed of pre- and post-synaptic compartments, and astrocyte peripheral processes.

To investigate whether sleep behavior is driven by one or multiple genes in this pathway, we examined whether these genes are differentially expressed in parental CC strains and functionally tested the roles of GRM5, GCPII, and GRM3 in sleep regulation using pharmacology. To assess strain-specific differences in gene expression, we performed qPCR on brain tissue from CC036 and CC057 mice. GCPII mRNA levels were significantly higher in the cortex, hippocampus, and hypothalamus of CC057 mice compared to CC036 (Fig.4D), whereas no differences were observed in mRNA expression for GRM3 or GRM5. Interestingly, Western blot analysis revealed significantly increased levels of mGluR3 and mGluR5 protein expression in the synapse-enriched membrane (P2) fraction from the cortex of CC057 mice (Fig.4E-F), suggesting a coordinated mechanism between increased GCPII transcription and glutamate receptor expression at the synapse.

To functionally test GRM3/5 and GCPII in sleep regulation, we treated parental CC036 and CC057 mice with the following pharmacological agents: GRM5 negative allosteric modulator MTEP (10mg/kg), GRM5 positive allosteric modulator LSN2463359 (LSN, 25mg/kg), GCPII inhibitor 2-PMPA (100mg/kg), and GRM2/3 inhibitor LY341495 (LY, 3mg/kg) (summarized in Fig.S4A). Since wake-promoting effects are expected to be more evident in the light phase when mice typically engage in more sleep, and sleep-promoting effects in the dark phase when mice are more active^44^, both periods were tested in consecutive weeks. Mice were injected at either ZT0 or ZT12 with vehicle control and 72hrs later by drug, and the drug effects were compared to control using within-mouse comparison (see Fig.S4B). Overall pharmacology results are summarized in Fig.S4C-H, and key findings presented in Fig.4G-I. MTEP was not found to have significant sleep modulating effects in CC036 and CC057 mice, whereas GRM5 activator LSN significantly reduced light-phase sleep amount (Fig.4G) and bout length (Fig.S4F) in both strains, and showed a trend to suppress dark phase sleep in CC057 (Fig.S4D and G). Similarly, inhibition of GRM2/3 with LY suppressed light-phase sleep in both strains (Fig.4F) and suppressed dark phase sleep in CC057 (Fig.S4D and G). LY also caused fragmentation of light-phase sleep bout length in CC036, but not CC057, perhaps because sleep is already fragmented in CC057 (Fig.1D). GCPII inhibition with 2-PMPA, thereby increasing levels of the GRM3 agonist NAAG, promoted dark-phase sleep (Fig.4I), and showed a trend to increase light phase sleep amount in both strains (Fig.S4C and E). Significant sleep modulating effects observed in these pharmacology experiments support our QTL identification of functionally important “sleep genes”. These results suggest that activation of GRM5 is wake-promoting, whereas activation of GRM3 via NAAG is sleep promoting, and the balance between these effects may be controlled by the enzymatic activity or expression of GCPII. Reminiscent of prior genetic discovery of the basis for sleep need identifying the synaptic gene Homer1a, our current unbiased genetic discovery related to daily sleep patterns identifies a glutamatergic signaling network coordinated across the tri-partite synapse (Fig.4J), supporting synapses as a cellular interface between circadian and sleep biology for the control of daily sleep patterns.

## Discussion

The daily patterns of sleep behavior are described by the long-standing 2-process model^3^: sleep timing is controlled by the CR (process C), and sleep depth/amount is a function of a poorly understood homeostatic sleep drive (process S). Initially understood as independent phenomena, emerging evidence suggests that processes S and C are functionally coupled; however, the molecular or cellular basis for this coupling is unknown^5,7^. The CR is a cellular phenomenon, based on the molecular clock acting within most cells^4^. Understanding of a cellular level of sleep need is a rapidly emerging area with strong indications that the need for sleep has a biochemical basis centered on synapse homeostasis^2,8,13-15^. Using genetically diverse mice, we conducted unbiased discovery and found a strong genetic basis for the daily consolidation of sleep patterns. Surprisingly, we identified a glutamatergic signaling network as a mediator of daily sleep patterns. This work provides support for the excitatory synapse as a cellular site for functional interaction between processes S and C.

Convergence of CC036/CC057 parental phenotypic traits in F1 hybrids (Fig.2) and divergence in F2 hybrids (Fig.3) indicate variant alleles in this population powerfully affect daily sleep behavior. QTL mapping identified a genomic region containing excitatory synaptic signaling molecules GCPII and GRM5. This QTL further implicates GRM3, as GCPII regulates the extracellular levels of a neuroactive peptide NAAG, a selective agonist for GRM3^27^. Quantification of mRNA expression showed upregulation of GCPII in CC057 compared to CC036, across cortex, hippocampus, and hypothalamus, with no change in GRM3/5 (Fig.4). Despite no change in transcript levels, CC057 mice expressed significantly higher levels of GRM3/5 protein in the synapse-enriched membrane fraction (P2) from the cortex (Fig.4). While the mechanism of GRM3/5 protein stabilization at synapses is not known, this result supports a coordinated biological function amongst these genes.

Pharmacology targeting GRM5, GRM3, or GCPII significantly modulated sleep behavior, supporting identification of functional “sleep genes” (Fig.4 and S4). GRM5 is a post-synaptic receptor previously implicated in sleep homeostasis in humans and rodents^38-41,45^. Consistent with prior work, activation of GRM5 with positive allosteric modulators was found to be strongly wake-promoting^39,41^. GRM3 is localized to neuronal pre-synapses and astrocytic processes^28,42,43^. Activation of pre-synaptic mGluRs, suppresses glutamate release^43^, potentially limiting the activity of post-synaptic GRM5. GCPII extracellular enzymatic activity degrades the selective GRM3 agonist NAAG^24^. Inhibition of GCPII was found to increase sleep, presumably through an accumulation of NAAG, suggesting that GRM3 promotes sleep. Conversely, inhibition of GRM3 with the non-selective GRM2/3 inhibitor (LY) was strongly sleep-suppressing, reinforcing the role of GRM3 as sleep-promoting. Collectively, these molecules form a coordinated glutamatergic signaling network acting across the excitatory tri-partite synapse. These findings support a model in which GRM5 and GRM3 promote wake or sleep respectively, and GCPII mediates biochemical switching between these pathways by regulating extracellular levels of neuroactive NAAG. (Fig.4J). Based on this model, broad upregulation of GCPII expression in CC057 mice may explain the highly fragmented sleep and loss of daily rhythm in this strain due to constitutive depletion of NAAG, subsequent loss of GRM3 activity, and increased GRM5 activation. Appropriate coordination amongst GRM5 and GCPII-GRM3 may enable the consolidation of wake and sleep behaviors to specific phases of the day, allowing for the highly consolidated sleep behavior seen in CC036.

### Excitatory synapses as a basis for the accumulation and resolution of sleep need

Sleep is a brain-wide state; the need for sleep is not exclusive to a certain brain region but is distributed across the nervous system. In a recent study, Weiss and colleagues used Drosophila to examine the brain-wide effects of sleep deprivation (SD) on neuronal synapses in a cell-type selective manner, revealing that excitatory neurons were almost exclusively responsive to SD, and that SD affected excitatory synapses uniformly across the entire brain^46^. The effect of SD in murine cortex has been examined using single cell RNA sequencing, showing that excitatory neurons are by far the most responsive cell type^47-49^. Isolation of forebrain synapses followed by “multi-omics” analysis revealed synapses undergo profound remodeling during the wake/sleep cycle involving the regulation of hundreds of proteins and thousands of phosphorylations^13,14,50-52^. This work further supports the synapse as a cellular interface between sleep and CR biology, where synapse localized mRNA are delivered to synapses based on circadian time of day, and synapse proteins and phosphorylations are regulated as a function of wake/sleep states^51,52^.

Cortical NREM-SWA is considered an established physiological marker for the accumulation and resolution of sleep need^2,5^. However, it is not clear how the dynamics of NREM-SWA reflects the molecular or cellular basis of sleep-need. Sawada et al., showed that the strength of excitatory synapses, specifically between excitatory neurons in the frontal cortex, but not inhibitory neurons, is a causal driver of NREM-SWA^53^. This is consistent with SHY which predicts the accumulation, and subsequent resolution of sleep need, is a function of broad excitatory synapse strengthening and weakening respectively^2^. In a seminal study, Franken et al., showed that the dynamics of NREM-SWA has a genetic basis and therefore a molecular substrate^8^. Subsequent efforts identified Homer1a as a molecular correlate of sleep need^9,10^. Homer1a drives homeostatic scaling-down of excitatory synapses through its interaction with GRM5^12^. Accordingly, we showed that Homer1a drives scaling-down during sleep^13^. It is remarkable that our current effort to examine the genetic basis of daily sleep patterns identified a QTL containing GRM5, the known target of Homer1a. Thus, independent, unbiased genetics efforts identify the Homer1a-GRM5 signaling pathway as a central mediator of sleep need and consolidated daily sleep patterns. Collectively, an emerging view supports excitatory neurons and excitatory synapses as a conserved and distributed locus for the accumulation and resolution of sleep need.

### Glutamatergic signaling coordinated across the tri-partite synapse

Astrocytes are rapidly emerging as important players in sleep regulation and function^54^. Astrocytes form functional components of synapses through interaction with neuronal pre-/post-synaptic compartments, collectively referred to as the “tri-partite synapse”^55^. Astrocytes occupy territories that tile much of the nervous system, form functional gap junctions, and are known to simultaneously interact with 100K-1M synapses^56^. Thus, astrocytes are well positioned to mediate coordinated synaptic processes during the wake-sleep cycle^54^. Astrocytes exhibit highly divergent patterns of intracellular Ca^++^ dynamics between wake and sleep states, suggesting astrocytes play an active and poorly understood role in sleep homeostasis and stability^57-59^. This builds on prior work demonstrating that “glial transmitters” play a key role in sleep homeostasis^54,60^. Furthermore, astrocytes sense surrounding neuronal activity through expression of GPCRs^61^. Using a chemogenetic approach, it was shown that Gi-coupled GPCR signaling in cortical astrocytes is sufficient to enhance sleep depth by increasing NREM-SWA^59^. Interestingly, GRM3 is Gi-coupled and is the primary mGluR expressed by adult astrocytes^28^. Our unbiased genetics discovery of GCPII-GRM3 signaling as a mediator of daily sleep patterns further implicates astrocyte synapse interactions in sleep function.

### Limitations

We utilized a non-invasive sleep scoring system PiezoSleep, which in this context enabled the throughput necessary to conduct our phenotypic screen and QTL mapping. By conducting continuous home-cage analysis over 5-7 days, this system provides robust measures of daily sleep patterns. However, unlike widely used EEG, this system does not provide definitive physiological metrics of sleep architecture or spectral power. During NREM-REM transitions, breath rate accelerates and becomes less regular, allowing for the estimation of NREM/REM using non-invasive methods^30,33^. Although this method is not yet fully established, it did allow us to collect estimated quantitative metrics for QTL mapping. We detected overlapping QTLs based on NREM, total sleep, and circadian metrics, perhaps reflecting that NREM makes up the majority of total sleep in mice^5^. We acknowledge that the lack of robust REM QTLs in this study may reflect a lack of suitable precision to accurately score REM metrics. Another possibility is that REM is highly polygenic and that our study lacked the statistical power needed to map relevant REM-QTLs, requiring a larger F2 population, and/or there was a lack of divergent alleles powerfully affecting REM sleep in parental CC036/CC057 strains. However, it is also likely that NREM and REM sleep have different underlying genetic and biochemical basis. In agreement, numerous advances in understanding the molecular basis of sleep regulation and function show that important wake/sleep regulating molecules such as PKA, calcineurin, SIK3, and HDAC4/5 have larger or exclusive effects on NREM, compared to REM^15,19-21,62^. Here, we base our conclusions primarily on the daily consolidation of sleep, based on multiple days of continuous recordings, which are readily enabled by the non-invasive methods utilized here.

### Conclusion

Sleep is a genetically complex trait^18^. Genetic mapping based on divergent daily sleep patterns in CC036 and CC057 mice identified GRM5 and GCPII-GRM3 as a coordinated glutamatergic signaling network affecting the daily consolidation of sleep patterns. Importantly, sleep and CR disruption are seen in many neuropsychiatric disorders^63^. GRM3/5 and GCPII are implicated in numerous psychiatric conditions, including schizophrenia, and have been linked with human cognitive ability, suggesting these are functionally important genes in human health and brain function^64-66^. Our unbiased genetic discovery of these pathways as regulators of daily sleep patterns may inform the mechanisms linking sleep/circadian disruption with brain health and psychiatric phenotypes.

## Materials and Methods

### Mice

All animal procedures were approved by the Institutional Animal Care and Use Committee of the University of North Carolina (UNC) at Chapel Hill and were performed in accordance with the guidelines of the U.S. National Institutes of Health. C57Bl/6J mice of both sexes were purchased from Jackson Labs. Collaborative Cross mice were obtained from the UNC Systems Genetics Core Facility (SGCF). The 13 Collaborative Cross strains included the following: CC011, CC013, CC027, CC033, CC036, CC037, CC041, CC057, CC058, CC060, CC071, CC072, CC081. CC036 and CC057 were selected for subsequent experiments. F1 hybrids were obtained from crossing CC036 females with CC057 males, and from crossing CC036 males with CC057 females, data was combined for F1 mice using both crosses described. F2 hybrids were obtained from inbreeding of F1 individuals. Experiments described in Figure 1 included males and females from all described strains. Subsequent figures utilized only female mice to mitigate anticipated issues with aggression. All experiments used adult mice aged 3-6months. Mice were maintained on a 12:12h light dark cycle with ad lib access to food and water.

### Sleep recordings and analysis

Sleep-wake behavior of adult C57BL/6J or CC mice was recorded continuously using a non-invasive piezoelectric monitoring system (PiezoSleep 2.0; Signal Solution, LLC, Lexington, KY). First, mice were transferred to our satellite facility containing PiezoSleep cages, maintained on a 12hr:12hr light:dark cycle. Mice were housed individually (15.5 cm^2^ cages) and provided with corn-cobb and cotton nestlet bedding material, crinkle paper and a wood chew for enrichment, water, and food. Following a 36-48hr acclimation period, sleep and wake behavior was recorded for 5-7 days for most recordings, uninterrupted, except for wellness checks. As described in pharmacology, some recordings lasted 2-weeks and included injections during continuous recording. No other experiments were conducted in the facility while recordings took place. Subjects were recorded in cohorts of 20-30 animals, and all equipment was thoroughly sanitized between recordings. The PiezoSleep monitoring system utilizes a piezoelectric mat positioned underneath the mouse cage to detect vibrational movements, including pressure exerted on the sensor by the mouse’s thorax during breathing. A customized statistics software (SleepStats, Signal Solution, Lexington, KY) processes these signals through an algorithm, distinguishing distinct respiratory patterns associated with sleep and wake states. Sleep is typified by periodic (2-3 Hz), regular amplitude signal, a common respiratory feature during mouse sleep states. In contrast, wakefulness is identified by the absence of standard sleep signals and the presence of irregular, higher amplitude signals, typically observed during voluntary movement (including subtle movements seen in quite wake). Using a linear discriminant classifier algorithm, the piezoelectric signals were classified as “sleep” or “wake” in 2-second epochs. Sleep-wake thresholds were automatically calculated for each individual mouse. To exclude brief or ambiguous arousals, a sleep bout is initiated when a 30 sec epoch contains greater than 50% sleep and is terminated when a 30 sec epoch contains less than 50% sleep. Validation of this software has been conducted using a combination of electroencephalography, electromyography, and visual assessment ^30-32^. NREM-REM transitions have been shown to be accompanied by transient accelerations of breath rate and decrease in breath rate regularity. Breath rate/regularity changes during sleep detected by piezoelectric sensors allow for the estimation of NREM and REM episodes^30,33^. Estimated NREM/REM amounts and bout lengths were analyzed using SleepStats 4.0 (Signal Solution, Lexington, KY).

We extracted sleep metrics using SleepStats (version 4.0) in bin sizes of one hour (percent) and 12 hours (percent, bout duration) starting from the light cycle onset at ZT0 (Zeitgeber time). Subjects from the Collaborative Cross survey recordings, as well as the experimental founder strains (CC036 and CC057) and the F1 hybrid generation, were averaged for group values. For multi-day sleep traces, the percent sleep for every hour was averaged across groups and plotted as consecutive days of recording (3-5 days). To generate a 24-hour “typical day” trace, each subject’s hourly percent sleep was collapsed across the recording and then averaged for the group. Hourly percent sleep was also used to calculate latency to sleep. Here, latency to sleep is defined as the number of hours ZT elapsed before the subject reached its mean sleep percent for that light cycle. Latency was calculated for each individual for each day, then averaged by subject and averaged by group. Bout durations were reported in seconds and were analyzed as recording means or as percent composition of all bouts recorded for each subject. All statistical analysis was performed in Graphpad Prism and are reported as means +/-SEM except for box and whisker plots (representing the median, 25th and 75th percentiles, and total range). C57Bl/6J was compared to CC strains using unpaired Student’s t-test with Bonferroni correction. CC036/CC057 parental strains, and F1 hybrids were compared by one-way ANOVA or multiple unpaired t-tests with Tukey’s correction for multiple comparisons, or 2-way ANOVA with Tukey’s multiple comparison test, as noted in figure legends. Light and dark cycle metrics were analyzed separately. For sleep recordings from the F2 generation all mice considered individually. Daily sleep metrics were calculated from F2 individuals by averaging 5-days of continuous recordings. Obtained metrics were used for QTL mapping.

### QTL mapping

We performed QTL analysis on an F2 mouse population generated from a cross between Collaborative Cross (CC) strains CC036 and CC057. Genotype sequencing was conducted on 13 mice from CC036 and 12 from CC057 to identify informative genetic markers. Single nucleotide polymorphisms (SNPs) were filtered to retain only those that were homozygous in both parental strains and had at most two missing genotypes per group. To ensure informativeness for QTL mapping, we selected SNPs that differed between CC036 and CC057.

Genotyping was then performed on F2 progeny from the CC036 × CC057 cross. For each SNP, we determined the parental origin of alleles in the F2 individuals. After quality control, we retained 258 F2 mice and 2,463 SNP markers for downstream analysis. QTL mapping was conducted using the qtl package in R (version 4.2.1). A single-QTL genome scan was performed across all markers using non-parametric methods, an extension of the Kruskal-Wallis test. Genome-wide significance thresholds were determined empirically using 10,000 permutation tests, and loci surpassing the 5% genome-wide threshold were considered significant. We calculated the Spearman correlations between phenotypes. The size of each circle showed the magnitude of the correlation coefficient, with blue indicating positive and red indicating negative correlations.

### Pharmacology

Pharmacology agents were purchased from Cayman Chemical (Anne Arbor MI, USA). The following drugs, and final concentrations were used for pharmacology experiments: GRM5 positive allosteric modulator LSN2463359: 25 mg/kg; GRM5 negative allosteric modulator MTEP HCl: 10 mg/kg; GCPII inhibitor 2-PMPA: 100 mg/kg; GRM2/3 antagonist LY341495: 3 mg/kg.

Drugs were dissolved in DMSO, and then diluted with 0.9% saline containing TWEEN 80 to a final vehicle preparation containing 5%DMSO, 5%TWEEN 80, 90% saline. Collaborative cross mice (CC036 & CC057) were moved into the PiezoSleep home-cage recording system, maintained on a 12:12h light/dark cycle, and were acclimated to this environment for 2-3 days without intervention, followed by a two weeklong experiment in which each mouse received 4x 100μL vehicle/drug intraperitoneal (IP) injections, two at light onset and two at dark onset. In week one, once acclimated, each mouse received IP injection with vehicle at ZT0/7AM/ light onset. 72hrs later, mice received IP injection with their assigned drug at ZT0. At the end of one week in the recording cages, bedding material was replaced with clean material prior to the start of the second week of the experiment. Following 2 days of recovery from cage maintenance, mice received a second set of injections beginning with vehicle injection ZT12/7PM/dark-onset, and again 72h later with drug injection, the same drug that was injected in week one. 24hrs of sleep data was analyzed following each injection, drug effects were compared statistically to the vehicle day using a within mouse comparison.

### Brain tissue isolation and prep

Mice were briefly anaesthetized using isolflurane and decapitated at ZT4. Brain regions of interest including cortex, hippocampus and hypothalamus were dissected in ice-cold phosphate buffered saline, frozen on dry ice, and then stored at -80oC for subsequent analysis. Mouse brain tissue suspended in ice-cold homogenization buffer (320 mM sucrose, 10 mM HEPES, pH 7.4, 1 mM EDTA, 5 mM sodium pyrophosphate, 1 mM sodium orthovanadate, 200 nM okadaic acid) supplemented with protease inhibitor cocktail (Roche). Tissue was homogenized using a dounce homogenizer with 12–15 strokes on ice. Tissue homogenate was used for RNA extraction or further processed for Western blot. For whole tissue lysate preparation, an aliquot of homogenate was mixed with 5x RIPA buffer (750 mM NaCl, 125 mM Tris, 5% IGEPAL, 0.5% SDS, 2.5% sodium deoxycholate, 25 mM sodium pyrophosphate) and incubated on ice for 30 min. The mixture was centrifuged at 17,000 x g for 10 min. at 4°C, and the supernatant was collected. P2 synaptosomal fractions were prepared from the remaining homogenate and centrifuged at 1,000 x g for 10 min. at 4°C. The supernatant was centrifuged at 17,000 x g for 20 minutes at 4°C, and the pellet (P2) was resuspended in 1x RIPA buffer and incubated on ice for 30 min. After a final centrifugation at 17,000 x g for 10 minutes at 4°C, the supernatant was collected as the solubilized P2 fraction. Protein concentrations for whole lysate and P2 fractions were quantified using the Bradford assay (Bio-Rad) in triplicate. All samples were diluted to 1 μg/μL using 1x RIPA buffer and stored until use.

### Quantitative real-time PCR

Mouse brain tissue (cortex, hippocampus, and hypothalamus) was rapidly dissected and homogenized in ice-cold homogenization buffer (320 mM sucrose, 10 mM HEPES pH 7.4, 1 mM EDTA, 5 mM sodium pyrophosphate, 1 mM sodium orthovanadate, 200 nM okadaic acid) supplemented with protease inhibitor cocktail (Roche). Homogenization was performed with 12– 15 strokes using a dounce homogenizer for cortex or with a 1 mL syringe for hippocampus and hypothalamus. Total RNA was extracted using the RNeasy® Plus Mini Kit (Qiagen). RNA quality and concentration were assessed using a NanoDrop Lite Spectrophotometer (Thermofisher). cDNA synthesis was performed using the High-Capacity cDNA Reverse Transcription Kit with RNase Inhibitor (Applied Biosystems, #4374966) with 2μg of RNA input per reaction. qPCR reactions were prepared using TaqMan™ Fast Advanced Master Mix (Applied Biosystems) and run on MicroAmp™ Fast 96-well reaction plates using TaqMan™ Gene Expression Assays specific for: GAPDH (Mm99999915_g1), FOLH1/GCPII (Mm00489655_m1), GRM5 (Mm01317984_m1), and GRM3 (Mm00725298_m1). Each reaction was run with a 20 μL final volume and in quadruplicate. Plates were centrifuged at 1200 rpm for 5 min. at 4°C before amplification on a QuantStudio 7 Flex Real-Time PCR System (Applied Biosystems). Gene expression levels were normalized to GAPDH and quantified using the comparative Ct (ΔΔCt) method.

### Western blot

Western blot samples were prepared by mixing protein lysates with 2x Laemmli sample buffer (20% glycerol, 100 mM Tris-HCl pH 6.8, 5% SDS, 50 mM DTT) and heating at 65°C for 15 min. Samples were then gently vortexed, centrifuged briefly, and placed on ice for 5 min. A total of 3 μg of protein per sample was resolved by SDS-PAGE on a 10% Tris/Glycine SDS-polyacrylamide gel and transferred to a nitrocellulose membrane (GE Healthcare). Membranes were blocked in 3% non-fat dry milk in 1x Tris-buffered saline (TBS; 50 mM Tris-HCl, 150 mM NaCl, pH 7.5) for 30 minutes at RT. Membranes were incubated overnight at 4°C with the following primary antibodies diluted in 3% BSA in 1x TBS with 0.05% Tween-20 and 0.05% sodium azide: anti-mGluR5 (1:5,000; Abcam, ab76316, rabbit), anti-mGluR3 (1:50,000; Abcam, ab166608, rabbit), and anti-GAPDH (1:1,000; Cell Signaling Technology, #2118, rabbit). After incubation, membranes were 3x for 10 min in 1x TBST (TBS + 0.05% Tween-20), then incubated for 1 hr. at RT with appropriate IRDye-conjugated secondary antibodies (1:20,000, LI-COR Biosciences, goat anti-rabbit 800CW) diluted in 3% non-fat dry milk in TBST. Membranes were washed 3x for 10 min. in 1xx TBST and imaged using the LI-COR Odyssey CLx imaging system. Band intensities were quantified using LI-COR Image Studio software.

## Acknowledgments

The authors wish to acknowledge technical assistance from Bradley Allf and Shenée C. Martin.

## Figures and Tables

**Fig. S1.**
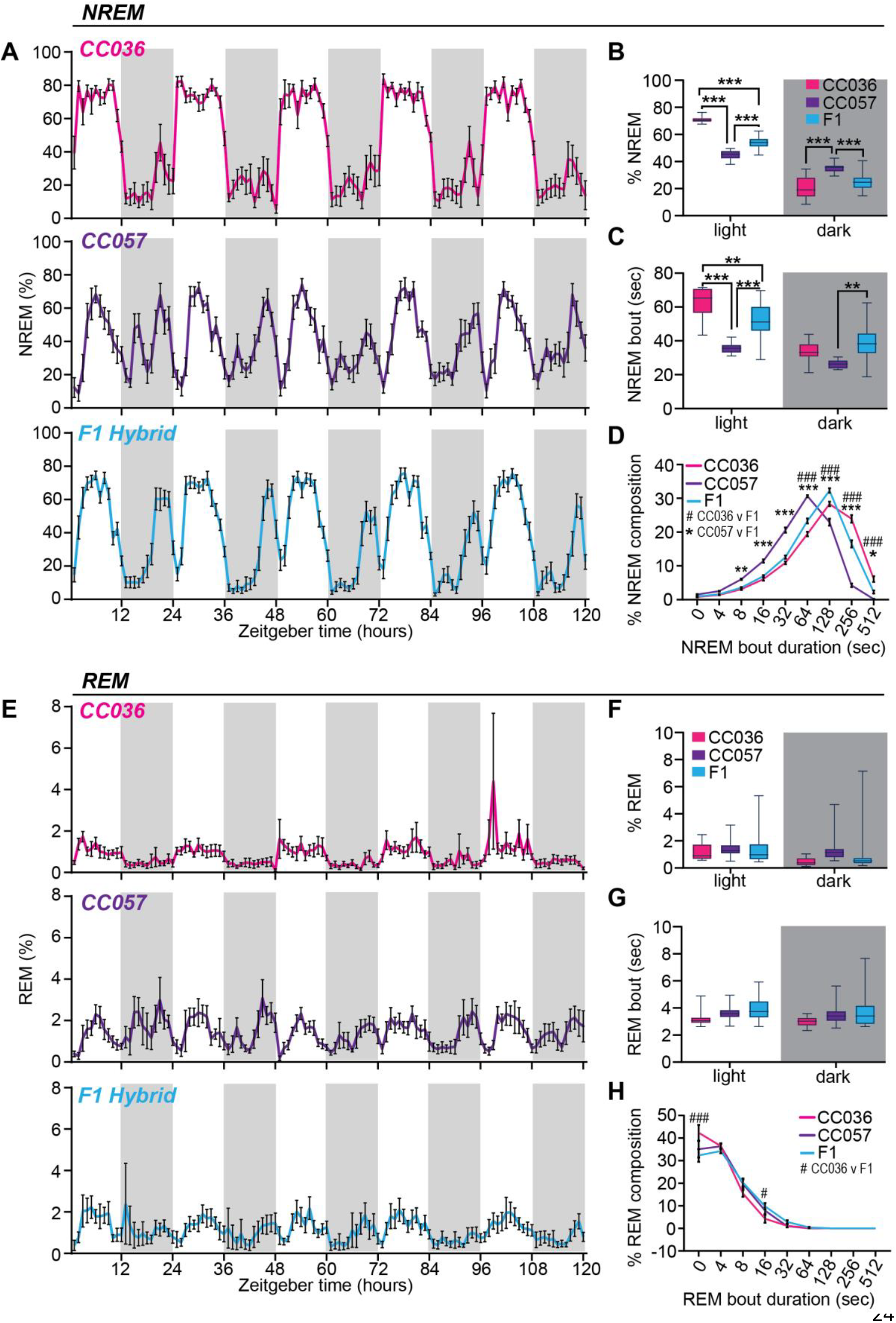
Convergence of CC036 and CC057 NREM sleep traits in the F1 hybrid and not REM traits. **A-D**. NREM sleep metrics estimated from breath rate analysis in female CC036 and CC057 parental strains and in the F1 hybrids. Error bars indicate standard error, N=10 females each for CC036 and CC057, and N=20 for F1 females. **A**. 5day traces of hourly % NREM sleep. Dark phase indicated by shaded area. **B-C**. quantification of average % NREM sleep (B) or NREM bout length (sec)(C) in light and dark phases. **P<0.01, ***P<0.001 1-way ANOVA with Tukey’s multiple comparison test. F1 hybrids show intermediate phenotypes in most NREM metrics between parental CC036 and CC057. **D**. Cumulative histogram of daily distribution of NREM sleep into bout lengths of varied duration. # and * indicates statistical difference between CC036 vs F1, and CC057 vs F1 respectively. *P<0.05, **P<0.01, ***/###P<0.001 2-way ANOVA with Tukey’s multiple comparison test. **E-H**. REM sleep metrics estimated from breath rate analysis in female CC036 and CC057 parental strains and in the F1 hybrids. **E**. 5-day trace of hourly %REM. **F**. Average % REM in light and dark phases. **G**. Average REM bout lengths (sec) in light and dark phases. No significant differences detected between groups, 1-way ANOVA with Tukey’s multiple comparison test. **H**. Cumulative histogram of daily distribution of REM sleep into bout lengths of varied duration. # indicates statistical difference between CC036 vs F1. #P<0.05, ###P<0.001 2-way ANOVA with Tukey’s multiple comparison test.

**Fig. S2.**
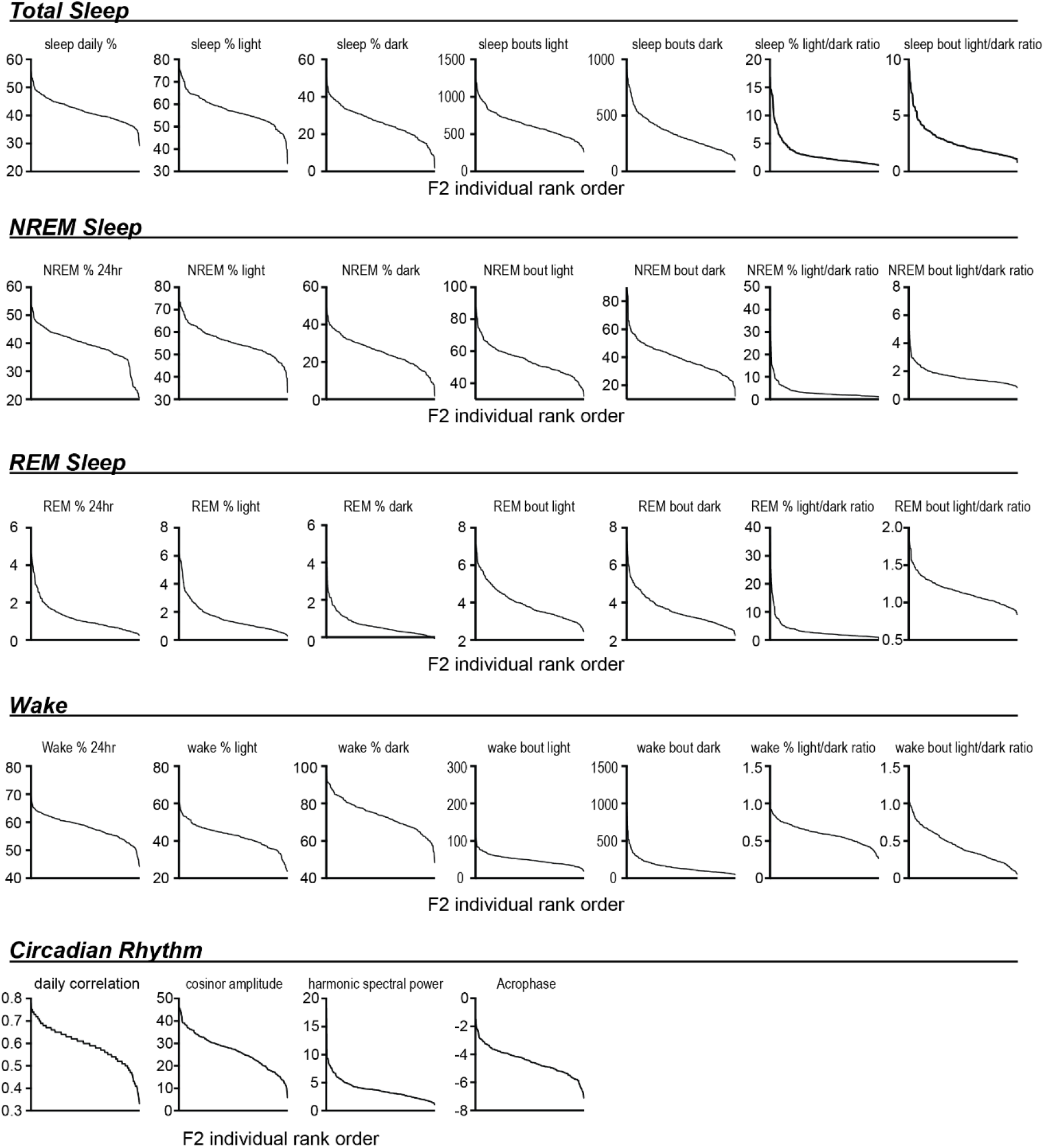
Summary of population data from F2 hybrids. 32 metrics of sleep and circadian behavior were measured from 279 F2 hybrids. Values are shown ranked highest to lowest to indicate the ranges of phenotypes across the F2 population.

**Fig. S3.**
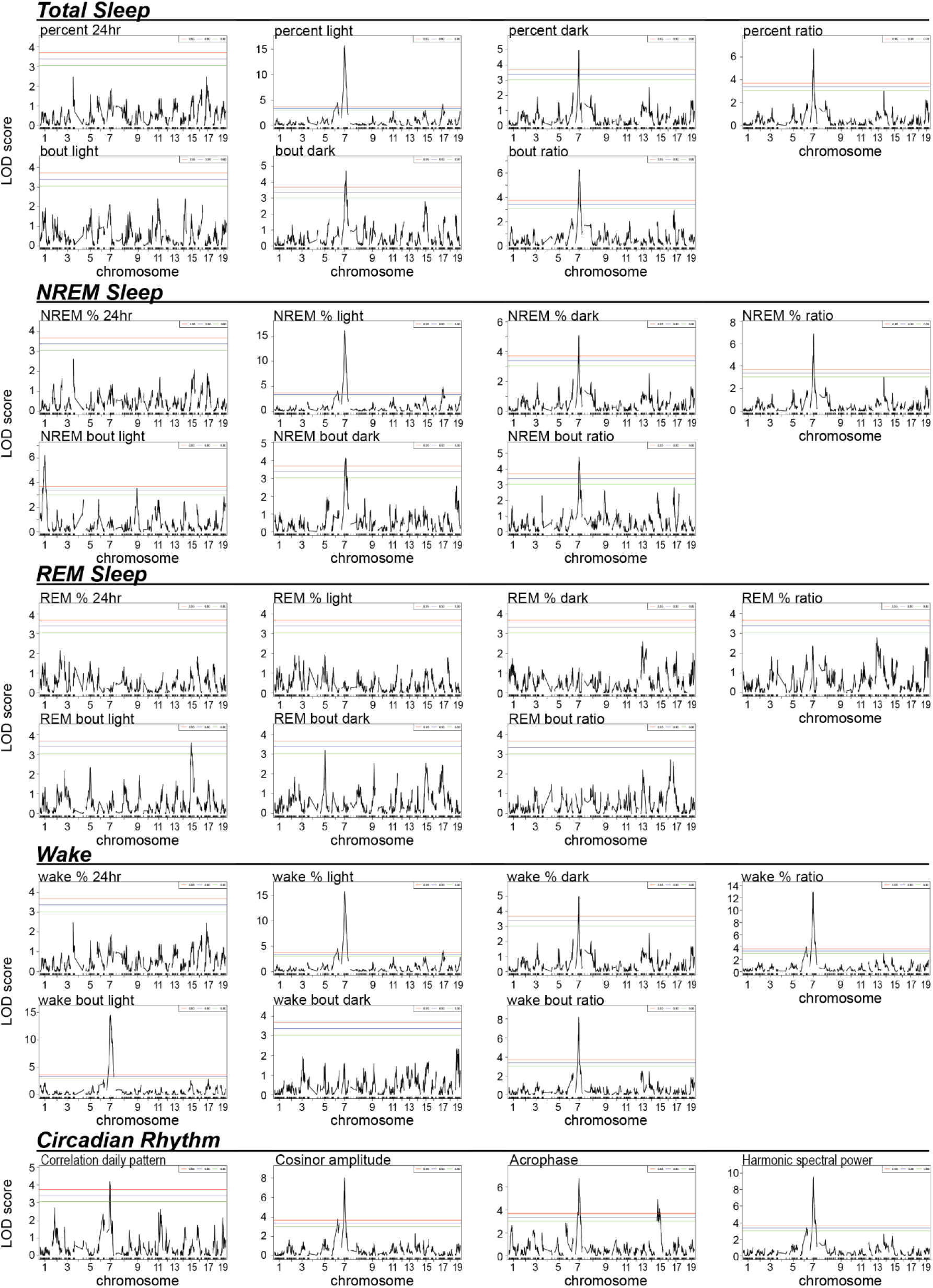
Summary of QTL analysis across 32 sleep and circadian metrics. Plots show genome-wide QTL analysis across 32 sleep and circadian metrics. Note the significant QTLs, above the 95% significance level, localized to chromosome 7 for 19 of 32 metrics. No significant QTLs, above 95%, were detected for REM metrics.

**Fig. S4.**
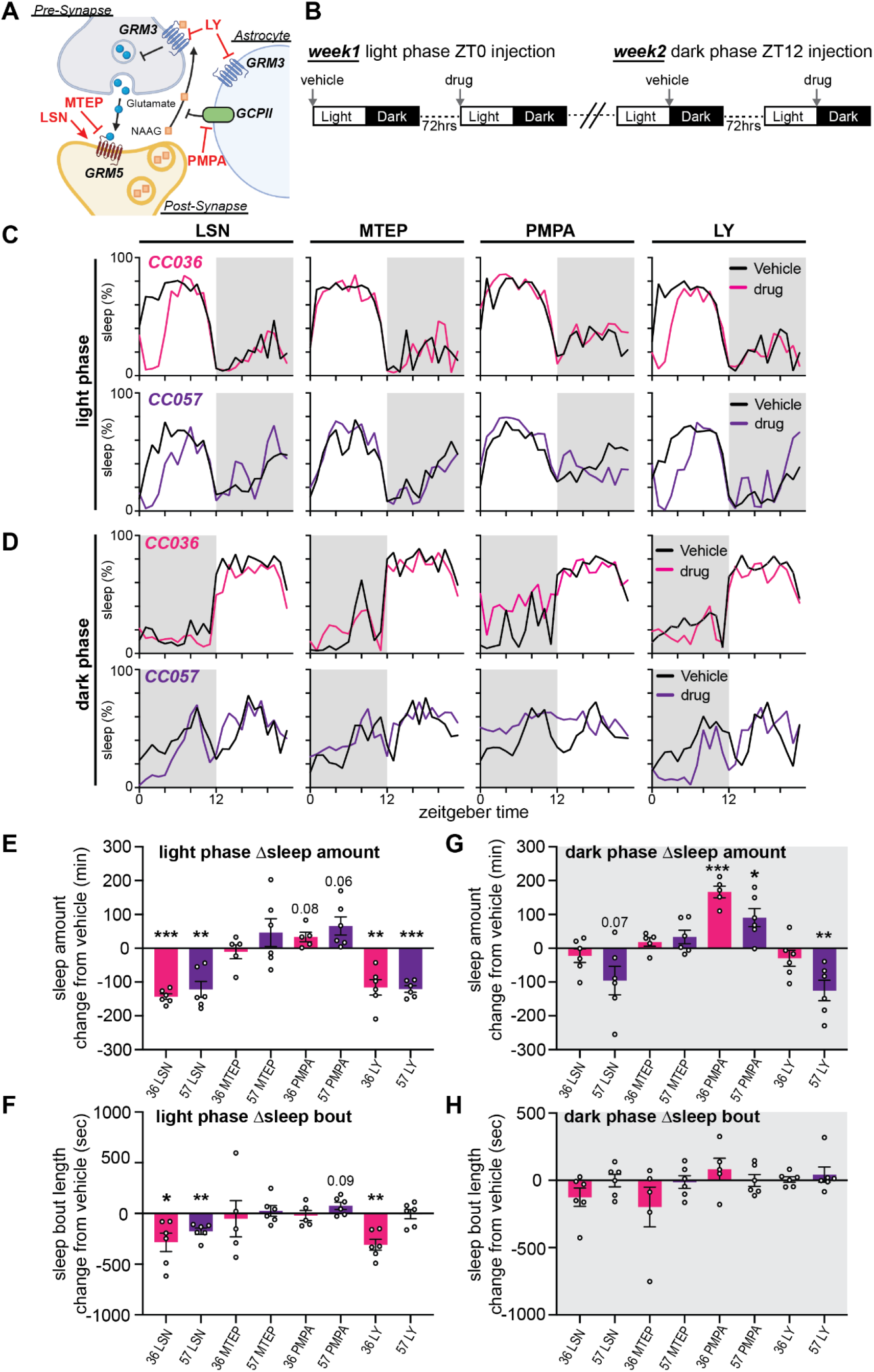
Summary of pharmacology validation. **A**. Model depicting coordinated signaling of GRM5, GCPII and GRM3 across the tripartite synapse, drugs targeting GRM5, GRM3, and GCPII are indicated, activator drug (LSN) indicated by red arrow, antagonists indicated by red plunger. **B**. Schematic of experimental design. In week1 of the experiment, CC036 and CC057 mice received vehicle injection, followed 72hrs later by drug injection at the beginning of the light phase, ZT0. In week2, mice received vehicle and subsequent drug injection immediately prior to the dark phase, ZT12. Sleep metrics following drug treatment were compared to the preceding vehicle treatment, using within mouse comparison. **C-D**. 24hr % hourly sleep traces following vehicle or drug injections at the beginning of the light phase (C) or immediately prior to dark phase (D) are shown. Traces are averaged from 5-6 individuals from each of CC036 and CC057 strains. **E-F**. Summarized effects of light phase injections depicted as change (Δ) in drug induced sleep amount (min)(E) or bout length (sec)(F) from drug injection compared to within mouse vehicle control. **G-H**. Summarized effects of dark phase injections in total sleep (min)(G) or bout length (sec)(H). N=5-6 mice per strain and treatment. *P<0.05, **P<0.01, ***P<0.001 Paired Student’s t-test.

**Table S1.** Summary of genome-wide significant QTLs. Table summarized significant QTLs located on Chromosomes 1, 6, 7, 9, 15, and 17. For each quantitative phenotypic trait the table lists statistical significance, logarithm of odds (LOD) score, chromosomal coordinates, upper and lower bounds, the number of genes and pseudogenes within the region, heritability score, and relevant SNPs flanking the region used for mapping. A minimum overlap amongst these QTLs is a 4.82MB region located on Chromosome 7 between coordinates 84.49-89.31MB.

| Phenotype | p-value | LOD | Chr | Position | Lower boi | Upper boi | Interval | snp | Heritability | gene | pseudoge | snp1 | snp2 |
| --- | --- | --- | --- | --- | --- | --- | --- | --- | --- | --- | --- | --- | --- |
| nREM_Bout_Light | 0.0001 | 6.43 | 1 | 107.97 | 76.7 | 135.82 | 59.11 | 23 | 11.13 | 927 | 235 | gbackupJAX00005508 | gUNC11739568 |
| Sleep_Percent_Light | 0.0072 | 4.64 | 6 | 114.39 | 113.3 | 127.89 | 14.59 | 15 | 8.15 | 423 | 91 | gUNC11882717 | SAC085115674 |
| nREM_Percent_Light | 0.0208 | 4.14 | 6 | 115.71 | 56.11 | 127.89 | 71.78 | 16 | 7.32 | 1444 | 442 | mUNC060379499 | SAC085115674 |
| Wake_Percent_Light | 0.0072 | 4.63 | 6 | 114.39 | 113.3 | 127.89 | 14.59 | 15 | 8.15 | 423 | 91 | gUNC11882717 | SAC085115674 |
| Wake_Percent_Ratio | 0.0122 | 4.4 | 6 | 119.19 | 113.3 | 127.89 | 14.59 | 15 | 7.75 | 423 | 91 | gUNC11882717 | SAC085115674 |
| Cosinor_Amplitude | 0.0225 | 4.13 | 6 | 119.19 | 113.3 | 142.69 | 29.39 | 27 | 7.29 | 750 | 224 | gUNC11882717 | gUNCHS019244 |
| Harmonic_Spectral_Power | 0.0354 | 3.87 | 6 | 119.19 | 113.3 | 142.69 | 29.39 | 27 | 6.85 | 750 | 224 | gUNC11882717 | gUNCHS019244 |
| Sleep_Percent_Light | 0 | 15.46 | 7 | 84.49 | 78.46 | 96.45 | 17.99 | 15 | 24.69 | 364 | 98 | gUNC13192480 | SS1073858155 |
| Sleep_Percent_Dark | 0.003 | 4.98 | 7 | 84.49 | 71.18 | 89.31 | 18.13 | 15 | 8.74 | 364 | 101 | gUNC13090495 | ST2073572206 |
| Sleep_Bout_Dark | 0.0035 | 4.72 | 7 | 116.53 | 82.27 | 122.74 | 40.47 | 31 | 8.3 | 1003 | 284 | S3C073290825 | gUNC13761022 |
| Sleep_Percent_Ratio | 0 | 6.82 | 7 | 84.49 | 78.46 | 89.31 | 10.84 | 10 | 11.77 | 266 | 77 | gUNC13192480 | ST2073572206 |
| Sleep_Bout_Ratio | 0.0001 | 6.88 | 7 | 107.61 | 84.49 | 112 | 27.51 | 16 | 11.87 | 720 | 240 | gUNC13271165 | gUNC13625638 |
| nREM_Percent_Light | 0 | 15.99 | 7 | 84.49 | 78.46 | 96.45 | 17.99 | 15 | 25.43 | 364 | 98 | gUNC13192480 | SS1073858155 |
| nREM_Percent_Dark | 0.002 | 5.11 | 7 | 82.27 | 71.18 | 91.14 | 19.96 | 17 | 8.95 | 407 | 105 | gUNC13090495 | gUNC13354082 |
| nREM_Percent_Ratio | 0 | 6.98 | 7 | 84.49 | 78.97 | 89.31 | 10.33 | 9 | 12.01 | 248 | 77 | SX1073158913 | ST2073572206 |
| nREM_Bout_Ratio | 0.0027 | 5.13 | 7 | 87.69 | 82.27 | 114.9 | 32.63 | 20 | 8.98 | 825 | 258 | S3C073290825 | SAJ074595854 |
| nREM_Bout_Dark | 0.0073 | 4.52 | 7 | 107.61 | 82.27 | 130.69 | 48.42 | 38 | 7.97 | 1275 | 318 | S3C073290825 | gUNC13858606 |
| Wake_Percent_Light | 0 | 15.45 | 7 | 84.49 | 78.46 | 96.45 | 17.99 | 15 | 24.68 | 364 | 98 | gUNC13192480 | SS1073858155 |
| Wake_Percent_Dark | 0.003 | 5.01 | 7 | 84.49 | 71.18 | 89.31 | 18.13 | 15 | 8.77 | 364 | 101 | gUNC13090495 | ST2073572206 |
| Wake_Percent_Ratio | 0 | 12.84 | 7 | 79.19 | 78.97 | 89.31 | 10.33 | 9 | 20.99 | 248 | 77 | SX1073158913 | ST2073572206 |
| Wake_Bout_Ratio | 0 | 7.93 | 7 | 79.19 | 78.46 | 89.31 | 10.84 | 10 | 13.54 | 266 | 77 | gUNC13192480 | ST2073572206 |
| Wake_Bout_Light | 0 | 14.02 | 7 | 96.45 | 78.97 | 107.61 | 28.64 | 15 | 22.68 | 782 | 228 | SX1073158913 | mbackupUNC070399178 |
| Correlation_Daily_Patterns | 0.0231 | 4.09 | 7 | 79.12 | 71.18 | 94.96 | 23.78 | 18 | 7.23 | 454 | 117 | gUNC13090495 | mUNC070396427 |
| Cosinor_Amplitude | 0 | 8.05 | 7 | 79.19 | 71.18 | 89.31 | 18.13 | 15 | 13.73 | 364 | 101 | gUNC13090495 | ST2073572206 |
| Acrophase | 0.0001 | 6.9 | 7 | 84.49 | 79.71 | 108.92 | 29.21 | 13 | 11.9 | 805 | 253 | gUNC13209247 | SS1074356963 |
| Harmonic_Spectral_Power | 0 | 9.46 | 7 | 87.69 | 78.97 | 89.31 | 10.33 | 9 | 15.94 | 248 | 77 | SX1073158913 | ST2073572206 |
| nREM_Bout_Light | 0.0398 | 3.81 | 9 | 70.29 | 41.6 | 87.44 | 45.84 | 20 | 6.76 | 1075 | 187 | S6R091664066 | gUNC16819132 |
| Acrophase | 0.0022 | 5 | 15 | 17.14 | 7.06 | 77.21 | 70.15 | 45 | 8.76 | 916 | 252 | gUNC24984364 | gJAX00083879 |
| Sleep_Percent_Light | 0.0075 | 4.6 | 17 | 68.49 | 57.72 | 86.1 | 28.38 | 27 | 8.09 | 386 | 125 | mbackupUNC170138236 | gUNC28521709 |
| nREM_Percent_Light | 0.0009 | 5.36 | 17 | 68.49 | 60.77 | 76.19 | 15.42 | 16 | 9.37 | 206 | 72 | gUNC28135094 | gUNC28369711 |
| Wake_Percent_Light | 0.0074 | 4.61 | 17 | 68.49 | 57.72 | 86.1 | 28.38 | 27 | 8.11 | 386 | 125 | mbackupUNC170138236 | gUNC28521709 |

